# Evolution of subgenomic RNA shapes dengue virus adaptation and epidemiological fitness

**DOI:** 10.1101/267922

**Authors:** Esteban Finol, Eng Eong Ooi

## Abstract

Genetic changes in the dengue virus (DENV) genome affects viral fitness both clinically and epidemiologically. Even in the 3’ untranslated region (3’UTR), mutations could impact the formation of subgenomic flaviviral RNA (sfRNA) and the specificity of sfRNA in inhibiting host proteins necessary for successful viral replication. Indeed, we have recently shown that mutations in the 3’UTR of DENV2 affected its ability to inhibit TRIM25 E3 ligase activity to reduce interferon (IFN) expression, which potentially contributed to the emergence of a new viral clade during the 1994 dengue epidemic in Puerto Rico. However, whether differences in 3’UTRs shaped DENV evolution on a larger scale remains incompletely understood. Herein, we combined RNA phylogeny with phylogenetics to gain insights on sfRNA evolution. We found that sfRNA structures are under purifying selection and highly conserved despite sequence divergence. Interestingly, only the second flaviviral Nuclease-resistant RNA (fNR2) structure of DENV-2 has undergone strong positive selection. Epidemiological reports also suggest that nucleotide substitutions in fNR2 may drive DENV-2 epidemiological fitness, possibly through sfRNA-protein interactions. Collectively, our findings indicate that 3’UTRs are important determinants of DENV fitness in human-mosquito cycles.

**Highlights:**

- Dengue viruses (DENV) preserve RNA elements in their 3’ untranslated region (UTR).
- Site-specific quantification of natural selection revealed positive selection on DENV2 sfRNA.
- Flaviviral nuclease-resistant RNA (fNR) structures in DENV 3’UTRs contribute to DENV speciation.
- A highly evolving fNR structure appears to increase DENV-2 epidemiological fitness.

## Introduction

Dengue virus (DENV) is the leading cause of mosquito-borne viral disease globally. An estimated 100 million cases of acute dengue occur annually, some of which develop into life-threatening severe dengue (Bhatt et al., 2013). DENV exists as four antigenically distinct but genetically related viruses (DENV1 to 4), all of which can cause the full spectrum of disease outcome. A tetravalent dengue vaccine has been licensed in several countries for use to prevent dengue. However, its protective efficacy varied across the four serotypes of DENV and long-term protection was only observed in older children with at least one episode of prior DENV infection (Hadinegoro et al., 2015). Thus, despite application of this vaccine and current approaches to vector control, DENV will likely continue to be a major public health challenge in the coming years.

Dengue is distributed throughout the tropics and is now encroaching into the subtropical regions of the world, causing frequent and recurrent epidemics (Messina et al., 2016). Whereas several of these epidemics were caused by fluctuations in the relative prevalence of the DENV serotypes in a background of low herd serotype-specific immunity, genetic differences in DENV also appears to play a distinct role in epidemic emergence (OhAinle et al., 2011). Indeed, we showed that the 3’ untranslated region (3’UTR) of DENV genome contributes to the epidemiological fitness of DENV, both through its interaction with human (Manokaran et al., 2015). Nucleotide substitutions in the 3’UTR of DENV2 strains resulted in increased sfRNA levels. This viral non-coding RNA binds to TRIM25 protein to inhibit its deubiquitylation (Manokaran et al., 2015); without TRIM25 E3 ligase activity, RIG-I signaling for type-I interferon (IFN) induction was repressed. Reduced type-I IFN response, at least in part, contributed to the increased viral spread of these strains in Puerto Rico in 1994. More recently, we have also shown that sfRNA from the same DENV2 isolated during the 1994 Puerto Rico outbreak also disrupts the antiviral response in the salivary gland of the *Aedes* mosquito vector (Pompon et al, 2017). Similarly, changes in the 3’UTR sequence that resulted in increased sfRNA production was observed in a new DENV2 clade in 2005 that resulted in a dengue epidemic in Nicaragua (OhAinle et al., 2011; Manokaran et al., 2015). Likewise, nucleotide composition in the 3’UTRs also differentiated dominant from weaker DENV strains in Myanmar, India and Sri Lanka (Myat Thu et al., 2005; Dash et al., 2015; Silva et al., 2008; respectively) although the structural consequences and impact of those substitutions on viral fitness have yet to be experimentally defined.

DENV 3’UTRs can be functionally segmented into three domains. The two more downstream domains possess RNA structures necessary for viral genome cyclization, viral RNA synthesis, translation and replication (Alvarez et al., 2005 and 2008). These structures have been termed small hairpin (sHP) and 3’ end stem-loop (3’SL) in domain III, and dumbbell (DB) 1 and 2 in domain II. Remarkably, a 30-nucleotide deletion (Δ30) in DB1 generated attenuated DENV-1, 3 and 4 but not DENV2 strains (Men et al., 1996; Durbin et al., 2001). Chimeric live attenuated DENV vaccines were designed to bear a Δ30 DENV4 3’UTR. They appear to be promising live vaccine candidates in clinical trials (Durbin et al., 2013). Vaccination with these vaccine candidates protected human volunteers against live DENV challenge infection (Kirkpatrick et al., 2016).

The proximal segment of the DENV 3’UTR contains RNA structures that are resistant to host nuclease activity, such as that of 5’-3’ exoribonuclease 1 (Xrn1), resulting in the production of subgenomic flavivirus RNA (sfRNA) during infection (Piljman et al., 2008). These structures have been referred to with different names in the literature: ‘stem-loops’ (Shurtleff et al., 2001, Piljman et al., 2008; Villordo et al., 2015) or ‘Xrn1 resistant RNA’ (Chapman et al., 2014A). However, crystal structures of homologous RNA sequences from related flaviviruses, Murray Valley Encephalitis virus (MVEV) (Chapman et al., 2014B) and Zika virus (ZIKV) (Akiyama et al., 2016), revealed a three-way junction RNA folding, rather than a SL structure. More importantly, these RNA structures halt diverse exoribonucleases to produce sfRNA, not only Xrn1 exoribonuclease (MacFadden A et al., 2018). Therefore, we refer herein to these RNA structures as flaviviral nuclease-resistant RNA (fNR) structures, as initially termed by Piljman et al. (2008), and to differentiate them from the unrelated nuclease-resistant RNA structures in dianthoviruses (Steckelberg AL et al., 2018).

As the sequence and hence the RNA structures in the 3’UTR of the DENV genome appear to be an important determinant of epidemiological fitness, it is possible that this part of the genome contributes to DENV evolution. Here we report the results of a detailed bioinformatic analysis that included all publicly accessible 3’UTR nucleotide sequences from DENV. We combined free energy minimization and sequence comparative analysis – also known as RNA phylogeny – to estimate secondary and tertiary RNA interactions in the 3’UTR of the four DENV types. Our RNA phylogenetic and natural selection analyses provide an evolutionary framework for further exploration into the molecular, epidemiological and clinical consequences of variations in the 3’UTR of dengue viruses.

## Results

### Sequence identity and RNA structures in the 3’ untranslated region of dengue viruses

A multiple nucleotide sequence alignment across many homologous non-coding RNA sequences can depict several features of their evolutionary trajectories. The ncRNA length, sequence identity and nucleotide composition can unveil insertion/deletion events and conserved GC rich functional RNA segments. Our analysis on the 3’ UTR of dengue viruses confirmed the existence of substantial differences in the nucleotide sequence composition and length across and within DENV serotypes as previously reported (Shurtleff et al., 2001; Proutski et al., 1999; Men et al., 1996). The 3’UTR of DENV1 serotype is the longest (Mode 465; range 436–475) followed by DENV2 (Mode 454; range 444–469), DENV3 (Mode 443; range 429–455), and DENV4 (Mode 387; range 387–407) **(Table S1A)**. Their distal segments (domains II and III) are highly conserved **(Figure 1A)**. In contrast, domain I of 3’UTR exhibits high genetic variability, including multiple insertions, deletions and point mutations. Indeed, its most proximal region – the hypervariable region (HVR) – depicted a significantly poor nucleotide conservation (Average identity < 89% in all serotypes, P<0.001) and significant adenine enrichment (p<0.05) **(Table S1B)**. This finding suggests a lack of folded RNA structures as RNA viruses tend to accumulate adenine in non-base-paired and structurally flexible regions (van Hemert et al., 2013; Keane et al., 2015). In comparison with the HVR, domain I also possesses a semi variable region with a high level of nucleotide conservation (Average Identity ≥ 96% in all serotypes) and significantly higher GC content **(Figure 1A and Table S1B),** suggesting the presence of conserved RNA structures across DENV serotypes. These conserved stretches are separated by small adenine-rich sequences, which may serve as spacer to facilitate the proper folding of functional RNA structures.

**Figure 1.**
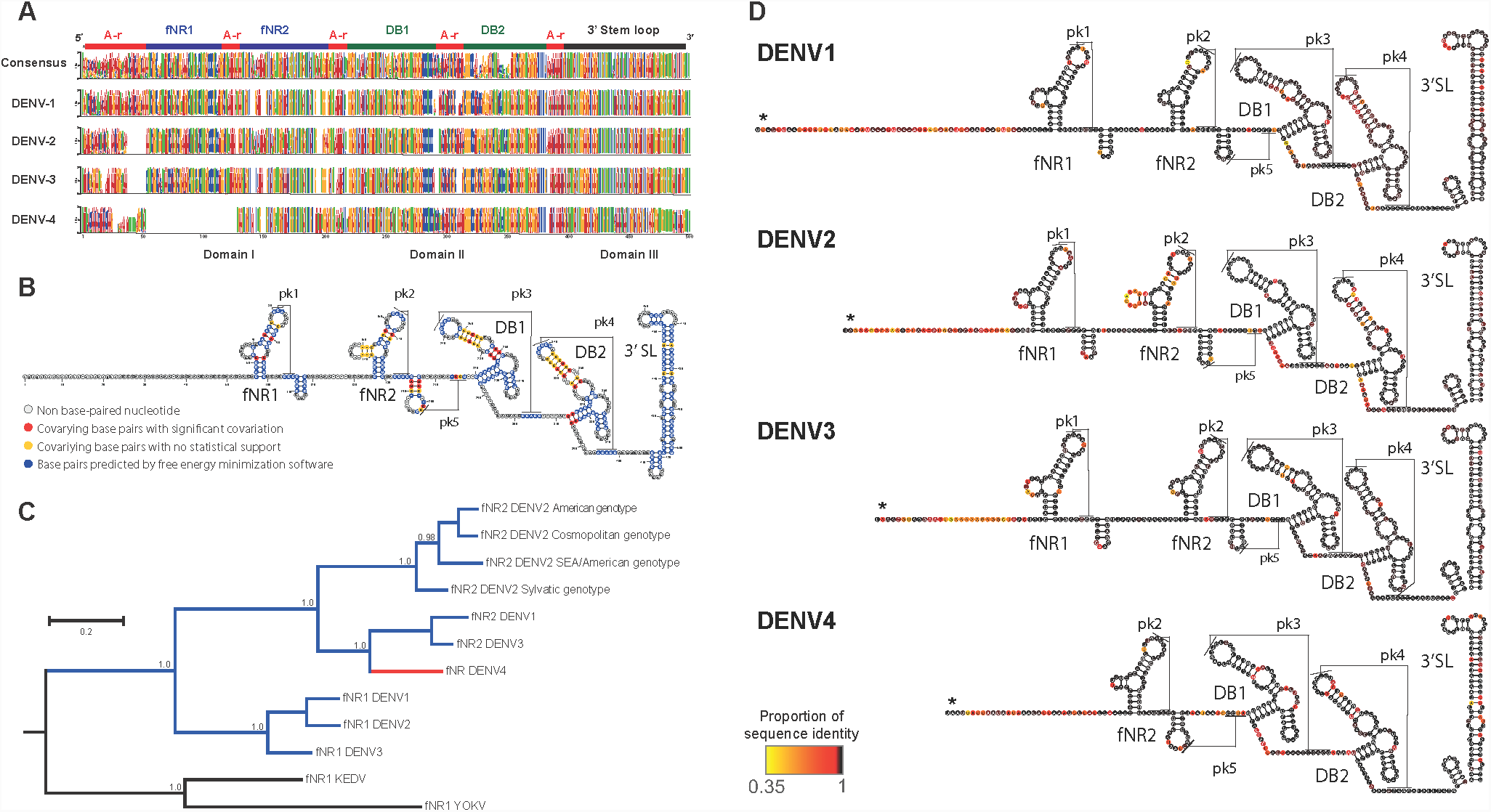
The 3’ UTR of dengue viruses diverged through deletion and sequence coevolution of functional RNA structures. Dengue viruses are phylogenetically related RNA viruses, their 3’UTR sequences have diverged along evolution, they now differ in sequence length and nucleotide composition. However, they kept functional RNA structures through sequence covariation. We observed and quantified RNA sequence covariation to predict secondary structures in the 3’UTRs and implemented Bayesian RNA phylogenetics to establish the phylogenetic relations among the RNA structures across dengue viruses. (A) Alignment of DENV 3’UTR sequence logos. Sequence logos for the 3’UTR of all DENV were aligned based on the multiple sequence alignment of DENV 3’UTRs sequences (consensus sequence logo). Highly conserved sequences were used to demark the boundaries between three domains in the 3’UTR of dengue viruses. Nucleotides are color-coded (Blue = Cytosine, Green = Uracil, Yellow = Guanine, Red = Adenine). Five conserved stretches were found across DENV 3’UTRs. They correspond to two flaviviral nuclease-resistant RNA (fNR) structures, two Dumbbell (DB) structures and the terminal 3’ Stem Loop (3’ SL). They were spaced by Adenylate rich (A-r) segments. This figure also illustrates the location of extra or missing nucleotides that account for the different lengths across 3’ UTR of DENV. The sequence logos also provide a glance on sequence conservation and nucleotide composition. These data are further described in Table S1. (B) Consensus model for the secondary structure of DENV 3’UTRs. After applying the RNA phylogeny approach, we obtained the secondary interactions for the five conserved RNA structures. Preliminary secondary structures and pseudoknots were predicted through free energy minimization and further refined by covarying base pairs. The statistical support for covarying base pairs was estimated by G-statistics in Rscape software (Figure S1 provides the parameters for the implementation and detailed results). (C) Phylogeny of fNR structures in the 3’UTR of dengue viruses. As the sequence logos revealed (A), DENV4 3’UTR bears only fNR structures, to determine whether this structure shares its most recent common ancestor with the fNR1 or fNR2 structures in other dengue viruses, Bayesian RNA phylogenetics was implemented under PHASE 3.0 software. It included all DENV fNR structure sequences (Branches in blue) and a DENV4 NR (Branch in red), the fNR structures from Kedougou virus (KEDV) and Yokose virus (YOKV) were used as outgroup (Branch in black). Posterior probabilities are only depicted on relevant nodes. (D) Ribonucleotide sequence identity on predicted RNA secondary structures in the 3’UTR of dengue viruses. Sequence identity is color-coded accordingly to the heat map at the bottom of the figure. Highly conserved sites are highlighted in a scale from red to black (Site conservation > 95%).

To define the secondary RNA structures and tertiary interactions of the 3’UTR, a ‘divide, learn and conquer’ approach was adopted. It included: (1) identification of conserved RNA structures within each DENV serotype; (2) prediction of preliminary RNA secondary structures from conserved short RNA segments using base-pairing probabilities and thermodynamic methods (Lorenz et al., 2011; Ren et al., 2005); and (3) validation, improvement and building of consensus RNA structures for DENV 3’UTR using RNA phylogeny. RNA phylogeny identifies evolutionarily conserved secondary and tertiary RNA structures through nucleotide sequence covariation (Jaeger et al., 1993). To further validate the covariation of base pairs, G-test statistics were also applied to determine whether these covariations occur at a rate higher than phylogenetically expected **(Figure S1)** (Rivas et al., 2017). The consensus DENV 3’UTR secondary structures derived from RNA phylogeny **(Figure 1B)** resembled the DENV2 3’UTR structure previously obtained by chemical probing (Chapman et al., 2014A), suggesting the validity of our bioinformatics approach. The predicted fNR structures in domain I are also compatible with the known crystal structures although they substantially differ from the secondary structures suggested by Shurtleff et al., 2001, and to a lesser extent, from the one reported by Villordo et al., 2015.

### fNR structures and DENV evolution

Interestingly, DENV1, 2 and 3 but not DENV4 bear two fNR structures **(Figures 1C and 1D)**. Among the duplicated fNR structures, the first fNR structure (fNR1) exhibited greater conservation than fNR2, suggesting that fNR1 exerts the main nuclease-resistant function for sfRNA production and is therefore necessarily highly conserved. The downstream fNR (fNR2) structure has thus relatively less constraint to evolve and adapt its structure possibly for additional function. The only fNR structure in DENV4 appeared to be phylogenetically related to the fNR2 than fNR1 structure of the other DENV serotypes, suggesting that the upstream stretch of nucleotides have been deleted **(Figure 1C)**.

To determine whether the variability of fNR structures influences DENV evolution, we examined the selection pressure on each nucleotide positions using a maximum-likelihood (ML) phylogenetic method (Wong and Nielsen, 2004). In this approach, the nucleotide substitution rate in each position of the sfRNA sequence was calculated and compared to the synonymous substitution rate in the coding region of each DENV serotype. This ratio models a ζ parameter. When a nucleotide position evolved neutrally, ζ ≈ 1 (i.e. equivalent substitutions rates). In contrast, ζ > 1 or ζ < 1 indicate that a given position in the sfRNA has respectively a higher or lower substitution rate than the synonymous substitution rate in the coding region of the genome from the same set of DENV strains. This approach thus provides an estimate on whether the higher substitution rate in a given nucleotide position of the ncRNA has contributed to increase the fitness of the bearing DENV strains, i.e. positive selection. If instead the ncRNA nucleotide position remains more conserved as compared to the neutral evolutionary rate of the coding region in DENV genome, it is regarded as the effect of purifying or negative selection. Figure 2A shows the ζ values for every position in the four DENV sfRNAs along with the 95% confidence intervals of the ζ parameter calculated for the coding region of the corresponding DENV genome (depicted as gray zone on the dot plots). Our results show that most nucleotide positions in DENV sfRNA have ζ<1, suggesting strong negative selection. This observation concurs with the predominant negative selection reported for the 3’UTR of DENV1 by Wong and Nielsen, 2004. It is also consistent with a reported finding that strong purifying selection characterizes the evolution of DENV genomes (Holmes, 2006 and Lequime et al., 2016). However, several nucleotide positions in the NR2 structure of DENV2 could have undergone positive selection with ζ>1 **(Figures 2A & 2B)**, suggesting that substitutions in fNR2 may confer competitive advantage for DENV-2 strains. These results are uniformly consonant with the greater dispersion and diversity of fitness peaks that appeared in our DENV sfRNA fitness landscape **(Figure 2C)**. The roughness on the surface of the DENV sfRNA fitness landscape indicates emergence and/or existence of multiple fNR/DB fitness peaks across the four DENV types and their divergence after fNR duplication. More importantly, the fNR2 genotypic space is populated by more dispersed and isolated peaks of fitness. It further confirmed the greater evolvability and divergences of fNR2 as compared to fNR1 sequences and the ability of fNR2 sequences to contribute to DENV type 2 divergence into different genotypes.

**Figure 2.**
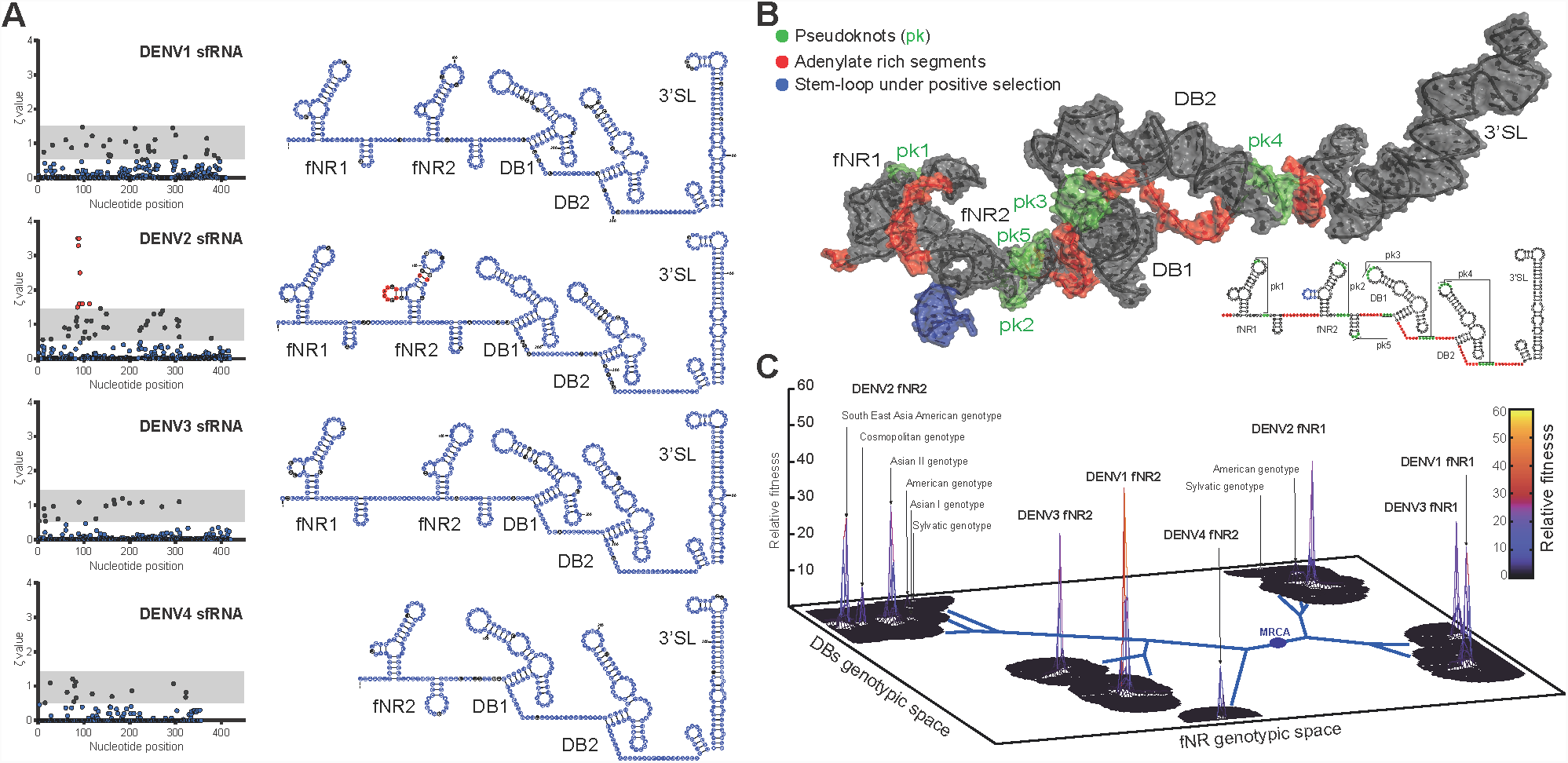
DENV2 sfRNA possesses a highly evolving RNA structure. (A) Site-specific quantification of natural selection on the sfRNA from dengue viruses. A maximum-likelihood method was applied to detect the action of natural selection in DENV sfRNAs in a site-by site basis. A zeta parameter and its 95% CI interval across the full genome determined whether a nucleotide position underwent negative (Blue dots), positive selection (Red dots) or neutral evolution (Black dots) in the ncRNA sequence. On the left, dot plots depict the Zeta values for all DENV sfRNAs. The 95 % CI is shown as gray zone on the dot plots. The 95 % CI slightly varied across DENV genomes. DENV1 = [0.513; 1.469], DENV2 = [0.547; 1.473], DENV3 = [0.532; 1.451], DENV4 = [0.506; 1.437]. On the right side, the ribonucleotide positions in the DENV sfRNA secondary structures are color-coded accordingly. Pseudoknots are not shown for the sake of simplicity. (B) 3D simulation of DENV2 sfRNA. By combining predicted base pairing *a*nd comparative RNA modeling, we obtained an *in-silico* 3D model of DENV2 sfRNA. Pseudoknots, adenylate rich regions and the highly evolving harpin are highlighted in colors as in the secondary RNA structures at the bottom right of the figure. (C) Fitness landscape of DENV sfRNA as determined by fNR and DB sequence abundance. A fitness landscape is a function that assigns to every genotype a numerical value proportional to its fitness. It involves a vast 2D genotypic space and a fitness value (Z axis). The 2D genotypic space for the sfRNA fitness landscape was resolved in X axis by the fNR nucleotide sequences and in the Y axis by the DBs (full length Domain II) sequences. whereas their normalized combined abundance (copy numbers) across DENV 3’UTR sequence alignments served as relative fitness value. It allowed us to elucidate the ability of these RNA structures to contribute to genome survival and to establish distinct evolutionary trajectories in sfRNA evolution.

To further assess the role of DENV2 3’UTR evolution, a ML phylogenetic tree was constructed using a nucleotide substitution model of evolution for non-coding RNA, based on the consensus secondary RNA structure of DENV2 sfRNA. The phylogenetic tree **(Figure 3A)** segregated the DENV2 sfRNA sequences into six clades, consistent with the six genotypes that characterize DENV2 evolution (Reviewed by Chen and Vasilakis, 2011).

**Figure 3.**
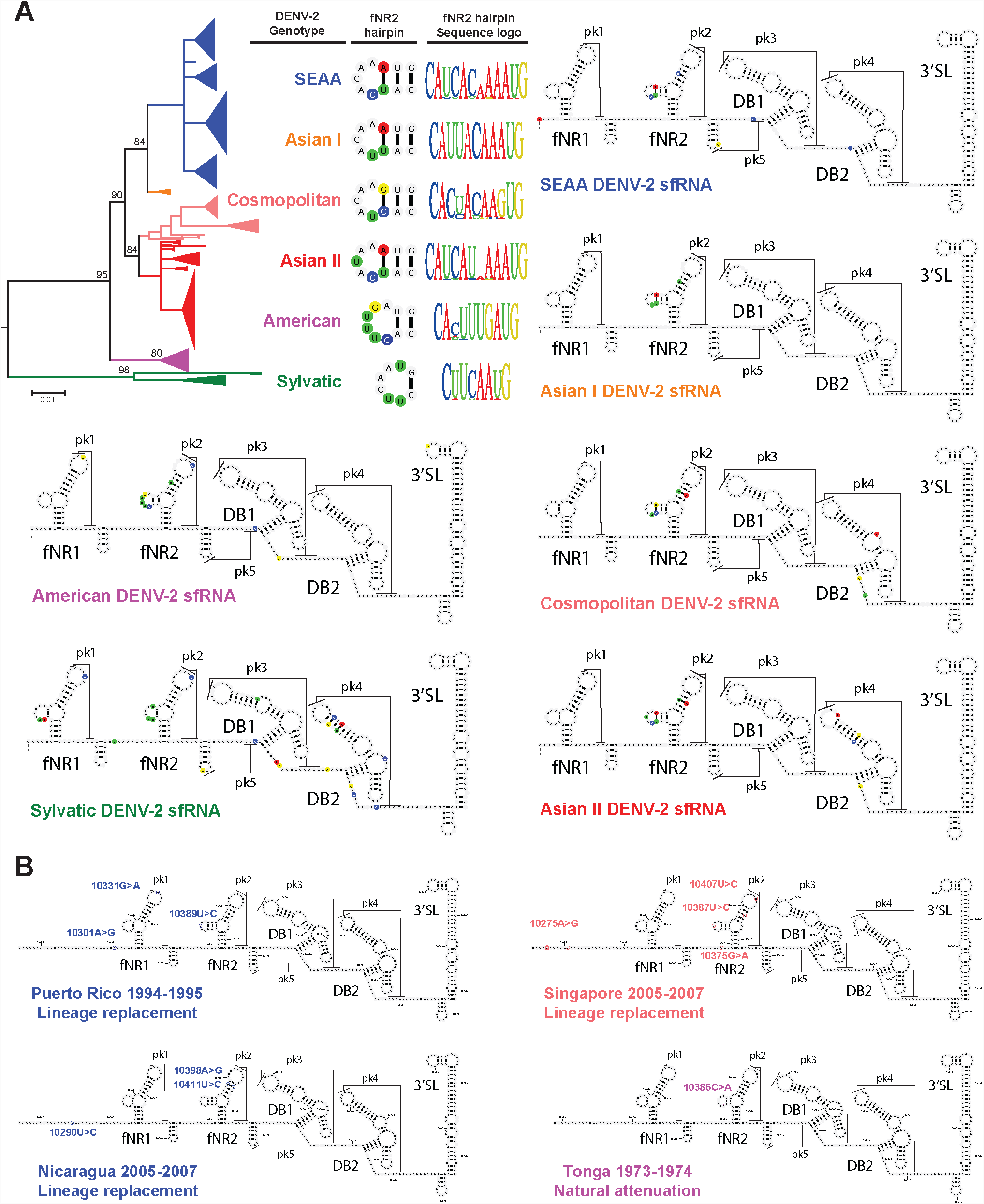
Nucleotide substitution in DENV2 sfRNA are associated to DENV2 speciation and increased epidemic potential. (A) Phylogenetics and nucleotide substitutions in the sfRNA of DENV2 strains. A maximum-likelihood phylogenetic tree from DENV2 sfRNA was built using PHASE 3.0 software. This software applies an RNA structure-based approach to construct phylogenies of non-coding RNAs. The highly evolving hairpin in DENV2 sfRNA exhibited distinct nucleotide composition and structure across DENV2 genotypes. Harpin secondary structures and sequence logos are shown next to the corresponding branch for the DENV2 genotypes in the phylogentic tree. The consensus secondary structures for the sfRNA in all the DENV2 genotypes is also shown. Genotype-specific nucleotide substitutions are highlighted in colors. (B) Epidemic DENV2 strains underwent nucleotide substitution in the highly evolving NR2 of DENV2 sfRNA. Location of nucleotide substitutions are shown in dominant strains that have been involved in three DENV-2 clade replacements and a natural attenuation event.

Remarkably, the positively selected hairpin in the fNR2 structure differed in nucleotide composition and structure across DENV2 genotypes, despite being relatively conserved within genotypes **(Figure 3A)**. Most genotypes are rich in adenine in this hairpin structure except the American genotype, which has mostly uracil. It is noteworthy that the American genotype has shown poor epidemiological fitness and has now been completely displaced by other DENV2 genotypes in many parts of the world. Likewise, analysis of DENVs derived from epidemiological studies **(Table 1)** also showed that four clade replacement episodes that resulted in greater or less than expected dengue incidence involved nucleotide substitutions in fNR2 structures **(Figure 3B)**.

**Table 1.** List of epidemiological events associated to increased DENV epidemiological fitness. An extensive literature revision on DENV epidemiology revealed at least 13 events associated with increased DENV epidemiological fitness. Nucleotide substitutions in the 3’UTR were reported in nine of those epidemic DENV strains (*).

| Events | Geographical location | Year | Serotype/Genotype | Context | reference |
| --- | --- | --- | --- | --- | --- |
| Clade replacement | Puerto Rico* | 1994 | DENV-2 SEAA | Outbreak | McElroy et al., 2011 |
|  | Nicaragua* | 2005 | DENV-2 SEAA | Outbreak | OhAinle et al., 2011 |
|  | Singapore* | 2005 | DENV-2 Cosmopolitan | Outbreak | Lee et al., 2012 |
|  | Thailand | 1990s | DENV-1 | Outbreaks | Teoh et al., 2013 |
|  | Brazil | 1990s | DENV-1 Genotype I | Outbreaks | Carneiro et al., 2013 |
|  | Mexico | 2006<br>1999 | DENV-1 Genotype III<br>DENV-2 SEAA | Several outbreaks | Carrillo-Valenzo et al., 2010 |
|  | India* | 2009-2011 | DENV-1 genotype III | Outbreaks | Dash et al., 2015 |
|  | Malaysia | 1987<br>1997<br>2004 | DENV-1 Genotype I | Recurring outbreaks | Dupont-Rouzeyrol et al., 2014 |
|  | Tonga Island* | 1974 | DENV-2 American | Reduced severity | Steel et al., 2010 |
| Genotype replacement | Sri Lanka* | 1989 | DENV-3<br>Genotype I> III | DHF emergence | Silva et al., 2008<br>Manakkadan et al., 2013 |
|  | Myanmar* | 1974-2002 | DENV-1<br>Genotype I> III | Several outbreaks | Myat Thu et al., 2005 |
|  | Indian Subcontinent* | 1971 | DENV-2<br>American>Cosmopolitan | Several outbreaks | Kumar et al., 2010 |
|  | Americas* | 1983 | DENV-2<br>American>SEAA | DHF emergence | Mir et al., 2014 |

## Discussion

The identification of 3’UTR structure and sfRNA production as having functional importance in determining viral fitness is of major interest in both experimental and epidemiological settings. The frequent emergence of DENV strains with insertions, deletions and point mutations in their 3’UTR (Zhuo et al., 2006, Pankhong et al., 2009, de Castro et al., 2013, Dash et al., 2015) and the differences in nucleotide lengths underscores the need for improved understanding of this part of the DENV genome. Given that the sfRNA is a non-coding RNA, its influence on DENV fitness and evolution must be understood in the context of its RNA structures. RNA phylogeny provides a bioinformatic approach to glean insights to direct further mechanistic investigations. Furthermore, a phylogenetic based estimation of substitution rate using non-coding RNA model of nucleotide substitutions coupled with normalizing by the substitution rate in the coding genome, enabled us to: (1) overcome the bias that dataset size can introduce in sequence identity (conservation) analysis; (2) avoid the misleading interpretation of “representative” sequences and; (3) exploit the growing sequencing databases to gain insight into the non-coding RNA evolution of a widely spread virus.

Available sequence data indicate that the DBs and the terminal 3’ SL (Domain II and III) are highly conserved. Their role in flaviviral replication may enforce a strong purifying selection on these 3’ UTR domains. The integrity of DB structures also contributes to increase sfRNA production, since the 30 bases nucleotide deletion in DB1 decreases sfRNA production and increases type I IFN susceptibility of the live attenuated DENV vaccine candidates (Bustos-Arriaga J et al., 2018). Likewise, fNR structures in the DENV 3’UTR are relatively well conserved. Any nucleotide substitution in base-paired positions of the fNR structures is often accompanied by compensatory mutation to maintain structural integrity. Along this line of thought, the finding of a positive selection in the NR2 structure of DENV 3’UTR is thus intriguing. fNR2 mutations may emerge during mosquito infection and subsequently be selected in human infections, due to their contribution to replicative fitness (Villordo et al., 2015; Filomatori et al., 2017). Indeed, serial passaging of DENV2 in *Aedes albopictus* derived C6/36 cells resulted in multiple mutations in its fNR2 and production of different sfRNA species (Villordo et al., 2015). These mutations could have then been positively selected in subsequent human infections in distinct geographical locations, possibly based on their ability to bind host proteins for the suppression antiviral immune activation. Indeed, Bidet et al. (2014) showed that fNR structure interacts with CAPRIN G3BP1 and G3BP2 proteins, mediating the sfRNA-induced repression of interferon-stimulated mRNAs in human liver-derived Huh7 cells. Moreover, mutations in the fNR2 structures produced higher replicative fitness in *Aedes albopictus* compared to the corresponding wild type DENV2 (Filomatori et al., 2017). Collectively, these findings suggest a strong evolutionary pressure on DENV2 fNR2 structures. Additionally, the consistent concordance between previously reported experimental data and our bioinformatics findings highlights the robustness of ML method developed by Wong and Nielsen (2004).

Given our RNA phylogeny findings and other available experimental evidence, we propose that the two fNR structures in domain I embody two functionally distinct RNA segments: the first NR structure is conserved to enable sfRNA production. In contrast, the downstream NR structure is relatively free to evolve and may be selected based on advantageous RNA-protein interactions in human or mosquito cells for increased fitness. This evolutionary model would agree with the reduced sfRNA production and transmission fitness that fNR1 mutations caused in DENV2 strains (Pompon et al. 2017), and the increased replicative, transmission and epidemiologic fitness that fNR2 mutations conferred to some DENV2 genotypes and more specifically in some dominant DENV2 strains. Indeed, Cologna and Rico-Hesse (2003) cloned the 3’UTR of the American genotype into a SE Asian/American DENV2 and found small viral plaques in Vero cells and slower growth kinetics in both mosquito and human cells, which are phenotypes more congruent with the American than SE Asian/American DENV2 genotype.

Our proposed model also explains the lack of positive selection in DENV4 as these viruses only possess one fNR structure. It, however, raises questions on why no positive selection was detected on the 3’UTR of either DENV1 or 3. We offer several interpretations. Firstly, If we assume that the distinct adaptation of the fNR2 in interspecies transmission – high sequence diversity in mosquito infection and sequence bottleneck in human infection – (Villordo et al. 2015 and Filomatori et al. 2017) is happening in all DENVs, our bioinformatics data would indicate that a stronger purifying selection occurs in the 3’UTR of DENV1 and 3 during human infection as compared to DENV2. A second and interesting scenario would suggest that the proposed model for the evolution of DENV2 sfRNA and its distinct fNR adaptation do not occur in other DENV types. This second postulate would help us understand why previous studies using RNA sequencing observed mutational hotspot in the 3’UTR of DENV2 but not DENV1 after replication in mosquitoes (Sessions et al. 2015; Sim et al. 2015). Experimental studies will be needed to test the validity of these postulates.

Notwithstanding the need for mechanistic validation, we suspect that fNR duplication has contributed to shape the overall divergence of dengue viruses. If the fNR duplication occurred early during DENV evolution – as the RNA phylogenetic analysis suggested, it is likely that the later fNR1 deletion in DENV4 imposed an evolutionary constraint in DENV4 lineage, limiting its adaptability to infect new ‘urbanized’ hosts and forcing a sympatric speciation and its greater divergence. This constraint might have been overcome through antibody-dependent enhancement in primates and/or competitive advantage in vector DENV co-infections in the current allopatric DENV distribution (Halstead, 2014; Vazeille et al., 2016). It would also help to explain why DENV4 has shown reduced epidemic potential during its global spread in the last decades and why only an additional 30 nucleotides deletion in the 3’UTR is required to generate a complete attenuated phenotype in DENV4 as well as in recombinant DENV1 to 3 strains bearing a Δ30rDENV4 -3’UTR (Durbin et al., 2001 and 2013).

Collectively, our findings suggest that 3’UTR evolution and sfRNA production are important determinants of DENV adaptation, survival and epidemiological fitness.

## Supporting information

## Acknowledgements

This work was supported by the Duke-NUS Signature Research Program funded by the Ministry of Health, Singapore and the Singapore National Medical Research Council. The authors would also like to thank Drs Vijaykrishnan Dhanasekaran and Mariano Garcia-Blanco for their invaluable advice throughout this work.

## Methods

### Sequence conservation analysis

Complete DENV genome nucleotide sequences were downloaded directly from the GeneBank database. The search included the keywords “Dengue virus type X” (X=1–4). In total, 1486, 1073, 831 and 154 sequences were included in the analysis of the 3’ UTR of DENV-1, DENV-2, DENV-3 and DENV-4, respectively. All DENV mutants, laboratory adapted strains, replicons, vaccine candidate strains, serially passage strains, and duplicated sequences were previously excluded. We built multiple sequence alignments for each serotype using MAFFT (Multiple sequence alignment using Fast Furier Transformation) software (Katoh and Standley. 2013). The sequence alignments were limited to the 3’UTR, starting from the Stop codon in NS5. We used Geneious platform to calculate nucleotide composition, sequence length (mode and range), average identity (i.e. average nucleotide conservation in the alignment) and number of identical sites (100 percent conserved positions) in the 3’ UTR of DENV (Kearse et al., 2012). To visualize the nucleotide composition pattern and conservation in the 3’ UTR of each serotype, we generated sequence logos from the 3’UTR alignments using Weblogo server (Crook et al., 2004) **(Figure 1A)**. We used standard colors to represent each type of nucleotide in the alignment (Blue = Cytosine, Green = Uracil, Yellow = Guanine, Red = Adenine).

To further characterize the nucleotide conservation, composition and distribution in the 3’UTR of dengue viruses, we identified some conserved stretches that mapped to the start and end of the Dumbbell (DB) structures in the Domain II, as described by Shurtleff et al. (2001) and Gerhald et al. (2011). We used them to establish a clear border between the different domains in the 3’ UTR of DENV and to perform subsequent sequence analyses in these domains. The statistical analysis of the nucleotide conservation and composition analysis was performed using STATA software (stataCorp, 2009). We used Analysis of Variance (ANOVA) with Bonferroni correction and Chi-square to test hypotheses from absolute values (average identity) and relative frequencies (nucleotide composition). Due to substantial differences across the sample size in the four data sets, all statistical comparisons were performed to test hypotheses within each serotype **(Table S1).**

### Sequence comparative analysis and RNA structure determination

To obtain secondary RNA structures and tertiary interactions, we applied a ‘divide, learn and conquer’ approach. It combines (1) an insightful 3’ UTR sequence conservation analysis within each flavivirus and within flavivirus groups to identify the presence of conserved RNA structures, (2) the power of RNA structure prediction software to solve preliminary RNA secondary structures from short RNA segments and (3) the robustness of a sequence comparative analysis – or RNA phylogeny – to validate, improve and build a consensus RNA structure for the 3’ UTR and sfRNA of flaviviruses (Jaeger et al., 1993). The strength of the RNA phylogeny approach relies upon the identification of evolutionary conserved functional RNA structures whose nucleotide sequences changed overtime but kept the RNA secondary and tertiary structures. Hence, it was possible to identify conserved functional RNA structures through sequence conservation analysis, to predict preliminary RNA structures using base-pairing probabilities and thermodynamic methods on the conserved stretches using RNAfold and HotKnots software (Lorenz et al., 2011 and Ren et al., 2005) and to validate secondary and tertiary interaction by identifying co-variations in the RNA nucleotide sequences, exploiting the growing sequencing dataset and high nucleotide substitution rate in RNA viruses. G-test statistics was implemented to further test whether observed RNA covariations occurred above phylogenetic expectation (Rivas et al., 2017) **(Figure S1)**. We drew secondary RNA structures and pseudoknots using VARNA software (Darty et al., 2009).

### Detecting natural selection in DENV sfRNA

To determine whether the RNA structures in the sfRNA play a role in the evolution of DENV, we explored natural selection pressure in a site-by-site basis in the sfRNA structure using a maximum-likelihood (ML) method (Wong & Nielsen. 2004) **(Figure 2A)**. We modeled the evolution of coding and non-coding regions and assumed a constant neutral (synonymous) nucleotide substitution rate in both regions in each serotype viral genome. We modeled the evolution in the open reading frame of DENV genome and determined its synonymous substitution rate, using a model of codon evolution (General Time Reversible, GTR+Γ) that has been generally applied to study the coding region of DENV genome (Weaver & Vasilakis. 2009). On the other hand, we calculated the nucleotide substitution rate in the sfRNA sequence in site-by-site basis. We used PHASE 3.0 software and a combined model of non-conding RNA evolution (Loop model: Hanley and Knott Regression, HKR+Γ and Stem model: 16D) based on the RNA secondary structure for the sfRNA from each serotype (Allen JE, Whelan 2014). We normalized the nucleotide substitution rate in each position of the sfRNA sequence by the synonymous substitution rate in the coding region of each DENV serotype and estimated a ζ parameter. Thus, a nucleotide position that exhibited a similar nucleotide substitution rate to the synonymous substitution rate (ζ ≈ 1) was assumed to be under a neutral evolution, whereas when ζ was found to be significantly higher or lower than 1 in a given position in the sfRNA sequence we assumed that it has experienced the action of positive or negative selection, respectively. To provide statistical significance to ζ parameter ratios, we calculated a 95% confidence interval (CI) for ζ parameter across each DENV serotype genome. If the ζ value of a given position was within the 95%CI, we confirmed neutral evolution. If the ζ value was above or below the 95%CI, we reported a significant positive and negative selection, respectively.

### DENV2 sfRNA Phylogenetics tree

We constructed a phylogentics tree for the sfRNA of DENV-2 from an alignment of 356 unique and representative 3’UTR DENV-2 sequences. We used PHASE 3.0 package (Allen JE, Whelan S. 2014) to build a maximum likelihood phylogenetic tree using the same composed model of nucleotide substitution (Loop model: HKR+Γ and Stem model: 16D) based on the predicted RNA secondary structures in the sfRNA of DENV-2. The statistical support for the topology of the tree was determined by 1000 bootstrap replications.

### DENV-2 sfRNA 3D modeling

We modeled the 3D RNA structure of DENV-2 sfRNA using RNA composer (Popenda et al. 2012). We used for the input file all the secondary and tertiary interactions that we obtained from the RNA phylogeny approach. The modeling of DENV-2 fNR structures was optimized through comparative RNA modeling, using ZIKV fNR crystal structure as template (5TPY). This was performed using ModeRNA software (Piatkowski et al. 2016). The local geometry in preliminary models were refined through energy minimization using the AMBER force field in the Molecular Modelling toolkit (Hinsen K. 2000). The final simulations were inspected for steric clashes using the *find-clashes* function in ModeRNA. The final sfRNA model was visualized, colored and labeled using pyMOL software **(Figure 2B)**.

### Construction of a sfRNA fitness landscape

A fitness landscape is a XYZ plot that depicts in the XY plane the genotypic space and every point in the genotypic space projects a fitness value to the Z plane. For the sfRNA fitness landscape, the extend of the genotypic space was defined in the X and Y planes by the product of all possible fNR and DB sequences. The number of possible sequences corresponded to sequence length to the power of 5, accounting for the possibility of having U, C, A, G or a missing nucleotide in every position. Thus, the overall genotypic space covers 1.3425^14^ points (fNR length = 75 nucleotides, BD length = 179 nucleotides, genotypic space = 75^5^x179^5^). Domain III of DENV 3’UTR was not integrated into the 2D genotypic space. Its length (102 nucleotides, Table S1) would substantially increase the size of the genotypic space (1.3425^14^x102^5^) and it will not contribute to resolve the 2D space due the high sequence identity in this part of DENV 3’UTR (>99%, Table S1). The fitness value was obtained from the relative combined abundance of fNR and DBs sequences among all DENV full genome sequences in NCBI database. Thus, if a combination of fNR-DB sequences conferred high fitness to the bearing strain, they would pass through subsequent viral generations in the mosquito-human interspecies transmission, they would be sampled and sequenced. The sequences would them appear in the NCBI database. Unfortunately, some strains have been more frequently sampled and sequenced than other strains and the NCBI database holds a sequencing bias. To adjust the fitness value and reduce the impact of the sequencing bias, a phylogenetic based normalization was implemented. Every clade was weighted according to it proportional contribution to the total number of nucleotide sequences from every DENV type, this weighting was then transferred to every sequence in the clade. Thus, if a clade contains 40% of all the DENV sequences – an over-sampled clade, every sequence in this clade would no longer count as 1 in the sum for the Z value, it will rather count as 0.6. If the clade represents 0.1% of the all sequences, every sequence in the clade will add 0.99 to the Z Value. FASTAptamer-cluster software was used to cluster sequences into sequence families based on a fixed Levenshtein edit distance (Alam KK, et al. 2015). The fitness landscape was plotted using gnuplot graphing utility in LINUX Operating System. An XYZ plot was generated by the *splot* command and the surface plot was obtained after a gaussian approximation to the XYZ raw data by the *gauss* kernel under *dgrid3d* command. Figure 2C shows only a small section of the overall fNR-DB genotypic space (3.38^-9^ %, 453882 XY points), where a Z value could be assigned. The rest of the genotypic space is completely deprived of fitness peaks and is not shown.

---

**Figure S1. G-statistics was applied to identify significant covariation of ribonucleotide base pairs in the 3’ UTR of dengue viruses.**

(A) Covariation scores survival functions. Two survival functions of scores are obtained upon G-test statistic implementation in R-scape software. In red, we drew the survival function of scores for all possible pairs in the input alignment excluding those proposed as base pairs. In blue, we plotted the survival function of scores for proposed base pairs in the input alignment. The survival function for the null alignments is depicted in black. The black line corresponds to fit to a truncated Gamma distribution of the tail of the null distribution.

(B) Summary of input alignment statistics, R-scape test parameters and R-scape output statistics.

(C) List of proposed base pairs with significant covariation. Nucleotide positions correspond to the position in the 2D model in figure 1B. Score and E-value are also shown.

**Table S1. A detailed analysis of RNA sequence alignments exposed the variability within and across DENV 3’UTRs.**

(A) Statistical analysis on the multiple sequence alignments from the 3’ UTR of dengue viruses. This table summarizes the sequence conservation analysis on this segment of DENV genome. It includes number of sequences, the mode and range of sequence length, the absolute and relative number of identical sites (100% conserved sites) and the average identity.

(B) Nucleotide composition and sequence conservation in distinct segments of 3’UTR DENVs. This table summarizes the conservation analysis of the different domains in the 3’ UTR of DENV. The comparison across the three domains within each dengue virus type revealed a significantly (p<0.05) higher CG content in domain II and a lower average identity in Domain I (p<0.05) (highlighted in red). Further analysis also revealed significant differences (highlighted in blue) between the Highly variable region (HVR) and the semivariable region (SVR) in Domain I.

