## Supplementary material for "Evolution of subgenomic RNA shapes dengue virus adaptation and epidemiological fitness"

## A

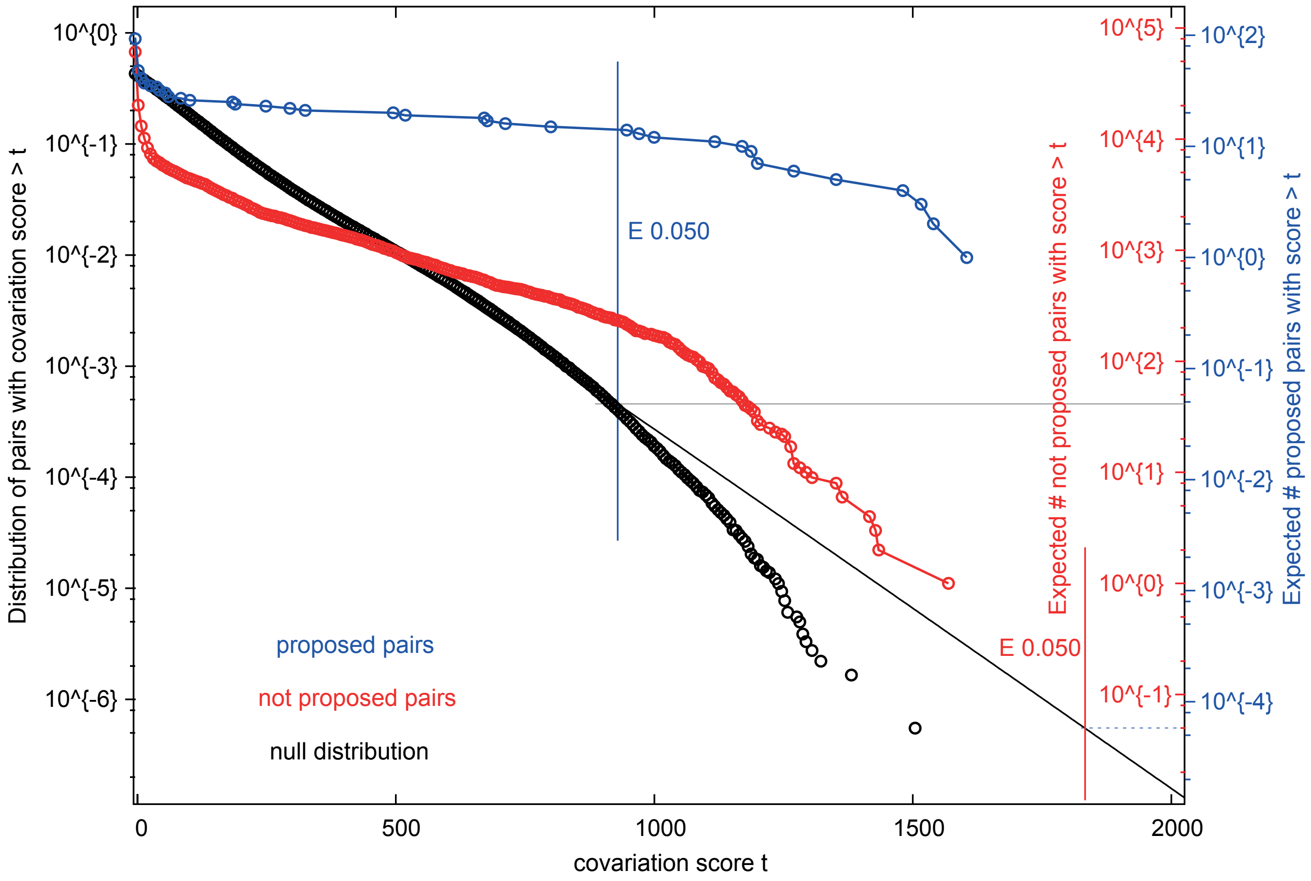


## B

**R-scape analysis:**

**Input file:**

- Multiple sequence alignment:
  - Number of sequences: 1825
  - Length: 501 nucleotides
  - Average identity: 83.17%
- RNA secondary structure:
  - Number of base pairs: 126

**R-scape test:**

- Covariation statistical method: APC-corrected G-Test statistic
- E-value threshold: 0.05

**R-scape output:**

- Number of base pairs after filter: 105
- Covarying base pairs: 14
- Covarying non base pairs: 0
- Range of scores: [-4.13 ; 1604.74]
- Sensitivity: 13.33
- Positive predictive value: 100

## C

**Base pairs with significant covariations**

| Left position | Right position | score | E-value |
|---|---|---|---|
| 61 | 95 | 973.31 | 0.0340113 |
| 74 | 90 | 1192.42 | 0.00682436 |
| 157 | 170 | 1486.24 | 0.00076987 |
| 188 | 205 | 1604.74 | 0.000307065 |
| 189 | 204 | 1355.32 | 0.00201342 |
| 199 | 215 | 1540.87 | 0.00049705 |
| 227 | 260 | 1204.55 | 0.00625533 |
| 228 | 259 | 1189.19 | 0.00682436 |
| 235 | 251 | 1275.27 | 0.00370847 |
| 313 | 384 | 1003.30 | 0.027391 |
| 314 | 383 | 1117.62 | 0.0115008 |
| 325 | 352 | 1174.07 | 0.00777609 |
| 329 | 348 | 947.33 | 0.0404361 |
| 331 | 346 | 1517.10 | 0.000592138 |
