## Supplementary material for "Evolution of subgenomic RNA shapes dengue virus adaptation and epidemiological fitness"

**A**

| Serotype | Sequences | Length | | Identical sites | | Average identity |
|---|---|---|---|---|---|---|
| | n | Mode | Range | n | (%) | (%) |
| **DENV-1** | 1486 | 465 | [436–475] | 169 | 35.4 | 96.5 |
| **DENV-2** | 1073 | 454 | [444–469] | 158 | 33.7 | 96.0 |
| **DENV-3** | 831 | 443 | [429–455] | 238 | 52.3 | 97.1 |
| **DENV-4** | 154 | 387 | [387–407] | 298 | 73.2 | 96.9 |

**B**

| 3' UTR Region | Serotype | Sequences | Length | | Nucleotide composition (%) | | | | | Identical sites | | Average identity |
|---|---|---|---|---|---|---|---|---|---|---|---|---|
| | | n | Mode | [Range] | A | G | C | U | GC | n | (%) | (%) |
| **Domain I** | DENV-1 | 1486 | 196 | [167–198] | 35.0 | 21.5 | 25.3 | 18.2 | 46.8 | 82 | (41.4) | **95.1** |
| | DENV-2 | 1073 | 183 | [173–184] | 36.7 | 21.9 | 22.8 | 18.6 | 43.7 | 66 | (35.1) | **93.4** |
| | DENV-3 | 831 | 175 | [161–180] | 30.4 | 24.6 | 27.8 | 17.2 | 52.4 | 91 | (50.0) | **94.8** |
| | DENV-4 | 154 | 113 | [113–131] | 31.3 | 26.9 | 21.4 | 20.4 | 48.3 | 76 | (58.0) | **93.4** |
| **HVR** | DENV-1 | 1486 | 51 | [22–51] | **55.8** | 13.6 | 14.4 | 16.2 | 28.0 | 1 | **(2.0)** | **88.1** |
| | DENV-2 | 1073 | 34 | [25–36] | **55.4** | 14.5 | 18.4 | 11.6 | 33.9 | 2 | **(5.6)** | **82.7** |
| | DENV-3 | 831 | 28 | [17–34] | **52.5** | 6.1 | 26.9 | 14.5 | 33.0 | 2 | **(5.9)** | **77.1** |
| | DENV-4 | 154 | 30 | [30–48] | **50.4** | 25.6 | 7.6 | 16.4 | 33.2 | 6 | **(12.5)** | **84.5** |
| **SVR** | DENV-1 | 1486 | 145 | [144–147] | 27.6 | 24.4 | 29.1 | 18.9 | **53.5** | 81 | (55.1) | 97.6 |
| | DENV-2 | 1073 | 151 | [148–154] | 28.0 | 23.5 | 28.8 | 20.2 | **51.3** | 65 | (42.2) | 95.7 |
| | DENV-3 | 831 | 147 | [147–148] | 26.5 | 27.9 | 28.0 | 17.6 | **55.9** | 89 | (60.1) | 98.1 |
| | DENV-4 | 154 | 83 | [83] | 24.2 | 27.3 | 26.7 | 21.8 | **54.0** | 70 | (84.3) | 96.8 |
| **Domain II** | DENV-1 | 1356 | 167 | [167–173] | 29.0 | 25.7 | 32.0 | 13.3 | **57.7** | 52 | (29.9) | 97.8 |
| | DENV-2 | 1071 | 169 | [169–171] | 31.6 | 24.7 | 29.9 | 13.8 | **54.6** | 75 | (43.4) | 97.6 |
| | DENV-3 | 822 | 166 | [166–169] | 28.7 | 26.8 | 31.5 | 13.0 | **58.3** | 94 | (55.6) | 98.8 |
| | DENV-4 | 153 | 172 | [172–174] | 29.7 | 26.6 | 31.3 | 12.4 | **57.9** | 138 | (58.0) | 98.4 |
| **Domain III** | DENV-1 | 1140 | 102 | [102–105] | 30.1 | 24.9 | 25.1 | 19.9 | 50.0 | 35 | (33.3) | 99.2 |
| | DENV-2 | 1008 | 102 | [102–103] | 30.7 | 24.8 | 24.1 | 20.4 | 48.9 | 30 | (28.8) | 99.5 |
| | DENV-3 | 715 | 102 | [102–104] | 29.1 | 24.7 | 25.4 | 20.8 | 50.1 | 53 | (51.0) | 99.6 |
| | DENV-4 | 151 | 102 | [102] | 28.8 | 24.6 | 25.9 | 20.7 | 50.5 | 84 | (82.4) | 98.5 |
